# Habitat type and interannual variation shape unique fungal pathogen communities on a California native bunchgrass

**DOI:** 10.1101/545632

**Authors:** Johannah E. Farner, Erin R. Spear, Erin A. Mordecai

**Affiliations:** Biology Department, Stanford University, Stanford, California 94305 USA; Smithsonian Tropical Research Institute, Panama City, Panama, Republic of Panama

**Keywords:** California grasslands, leaf spot pathogen, plant fungal pathogens, serpentine soil, Stipa pulchra

## Abstract

The role of infectious disease in regulating host populations is increasingly recognized, but how environmental conditions affect pathogen communities and infection levels remains poorly understood. Over three years, we compared foliar disease burden, fungal pathogen community composition, and foliar chemistry in the perennial bunchgrass *Stipa pulchra* occurring in adjacent serpentine and nonserpentine grassland habitats with distinct soil types and plant communities. We found that serpentine and nonserpentine *S. pulchra* experienced consistent, low disease pressure associated with distinct fungal pathogen communities with high interannual species turnover. Additionally, plant chemistry differed with habitat type. The results indicate that this species experiences minimal foliar disease associated with diverse fungal communities that are structured across landscapes by spatially and temporally variable conditions. Distinct fungal communities associated with different growing conditions may shield *S. pulchra* from large disease outbreaks, contributing to the low disease burden observed on this and other Mediterranean grassland species.

## Introduction

How environmental heterogeneity affects species interactions, and ultimately population growth rates and structures, remains a fundamental ecological question. Infectious disease, which arises from interactions between hosts, parasites, and the environment, is particularly likely to vary across populations in differing environments. Biotic factors, such as the phylogenetic structure of the host community, and abiotic factors, such as elevation, have been associated with the presence of specific diseases and the severity of their impacts on their hosts (Abbate and Antonovics 2014; Parker *et al*. 2015). Additionally, mounting evidence suggests that specific characteristics of pathogen community structures mediate the outcomes of host – pathogen interactions. The identities and abundances of the pathogen species present can shape infection dynamics both through pathogen – pathogen interactions, which can range from facilitative to antagonistic depending on the community context, and because pathogen species vary in their transmissibility, virulence, and sensitivity to different environmental conditions (Agrios, 2005; Seabloom *et al*. 2015; Borer *et al*. 2016; Dallas et al. 2019).

Separate observations of abiotic and biotic impacts on infection outcomes combine to highlight the importance of understanding interactions between environmental variation, pathogen community composition, and pathogen pressure on host populations. For plant hosts, identifying the conditions that place populations at high risk for severe disease outbreaks and applying preventative measures may prevent local species extinctions and improve crop yields. While studies have frequently explored the interactions of single host – pathogen pairs under differing abiotic conditions (e.g., Abbate and Antonovics 2014), both the effect of varying pathogen community composition on disease burden and the relationships between habitat characteristics and pathogen community structures remain largely unexplored in natural systems (but see Rottstock *et al*. 2014 and Spear 2017).

Fungal pathogens are particularly important to study in this context because they cause the majority of known plant diseases, have an increasing rate of emergence, and pose a substantial threat to plant health and population persistence in natural and agricultural settings (Agrios 2005; Fisher *et al*. 2012). Dramatic diebacks of North American chestnut trees and oaks caused by fungal pathogens (Anagnostakis 1987; Rizzo and Garbelotto 2003) and economically devastating crop losses caused by fungal rusts shared among wild and domesticated plants (Loehrer and Schaffrath 2011; Nazareno *et al*. 2018) demonstrate this threat. These examples indicate that understanding the communities of fungal pathogens that infect wild plants is important for developing effective protection strategies for both natural biodiversity and human food supplies. However, we lack basic information about the identities, diversity, and effects of the fungal pathogens that infect most plant species (Burdon and Laine 2019). Therefore, there is a pressing need for systematic surveys of these pathogen communities and their variation across natural habitats.

Evidence that abiotic and biotic factors drive both natural fungal endophyte community assembly in healthy leaves and fungal foliar disease dynamics in agricultural systems suggests that different habitats are likely to affect fungal foliar pathogen community composition and disease burden (Agrios 2005, Arnold 2007, Burdon and Laine 2019). For example, the repeated finding that foliar endophyte community structures change with temperature, humidity, and rainfall (Cordier *et al*. 2012, Zimmerman and Vitousek 2012, Bálint *et al*., 2015, Millberg *et al*. 2015, Giauque and Hawkes 2016, Barge *et al*., 2019) suggests that climate is likely to also shape fungal pathogen communities. Evidence from the plant pathology literature supports this expectation: for foliar fungal pathogens of wheat, barley, and lettuce, among others, infection dynamics are closely linked to temperature and moisture availability (Shaw 1986, Moss and Trevathan 1987, Eversmyer *et al*. 1988, Huber and Gillespie 1992, Magarey et al. 2005, Bernard *et al*. 2013, Clarkson *et al*. 2014, Velásquez *et al*. 2018). Timing and amount of precipitation can alter infection dynamics by washing away fungal spores and/or transmitting spores to new hosts (Madden 1997).

Additionally, local habitat characteristics, such as soil and plant physical and chemical properties, plant community composition, and plant genetics, can determine the availability and quality of susceptible and competent hosts for different fungal pathogen species (Snoeijers *et al*. 2000, Mitchell *et al*. 2002, Solomon *et al*. 2003, Bolton *et al*. 2008, Rottstock et al. 2014, Velásquez *et al*. 2018). Differences in soil nutrient content, plant nutrient content, and plant genetic backgrounds can influence the identities and abundances of fungal endophytes that inhabit the leaves of individual host species (Eschen *et al*. 2010, Busby *et al*. 2014). However, the short one- to two-year lengths of most observation periods hamper a more general understanding of the factors that shape foliar endophyte communities by generating snapshots of fungal community composition absent the context needed to determine whether observed patterns are broadly representative (but see Giauque and Hawkes 2016). Extensive work in agricultural systems wherein nitrogen fertilization and plant genetic diversity are both associated with disease prevalence and severity support the likely relevance of local habitat factors to fungal pathogens (Zhu *et al*. 2000, Solomon *et al*. 2003, Verosoglou *et al*. 2013).

Here, we examine the effects of habitat type on the burden and species composition of natural fungal plant pathogens infecting a single host species over a span of three years. Specifically, we take advantage of the natural occurrence of the California native perennial bunchgrass *Stipa pulchra* in two unique, adjacent grassland types to understand the extent to which habitat type and time influence the landscape of disease that a single plant species encounters, including foliar fungal pathogen community composition and disease severity. Stark differences in soil properties and plant community structures between serpentine and nonserpentine greenstone (hereafter, nonserpentine) grassland habitats result in a suite of environmental differences that may influence fungal pathogen infection, reproduction, and persistence (Harrison and Viers 2007; Burdon and Laine 2019). Additionally, high interannual variation in climate conditions and plant community composition in California grasslands may lead to temporal variation in infection dynamics through impacts on fungal growth and survival (Hobbs and Mooney 1991, Fernandez-Going *et al*. 2012).

We hypothesize that both disease burden and pathogen species composition differ on *S. pulchra* individuals across habitat types and years. In turn, we hypothesize that habitat type influences *S. pulchra* foliar tissue chemistry in ways that may affect plant – fungal pathogen interactions, with lower C and N content in serpentine plants growing in nutrient-poor soils. If so, these foliar chemistry differences may translate into differences in fungal community composition and either (a) higher disease burden in serpentine plants relative to nonserpentine plants, facilitated by negative impacts of limited plant resource availability on plant health (as in Springer *et al*., 2006), or (b) lower disease burden in serpentine plants due to lower plant nutrient content limiting resources available for fungal growth. By contributing novel information about previously uncharacterized plant fungal pathogen communities in California grasslands, the results of this study will provide new insight into the diversity and natural history of the fungi that infect the leaves of wild grasses in temperate climates. Further, the results of this study will aid in assessing disease risk for plant populations in natural and agricultural contexts by providing information about the spatial and temporal scales over which environmental heterogeneity can affect the species composition and severity of foliar pathogen infection.

## Materials and Methods

### Study system

All surveys and sample collection took place at Jasper Ridge Biological Preserve (JRBP), a 485-hectare site in San Mateo County, CA managed by Stanford University (37.4° N, 122.2° W). JRBP has a Mediterranean climate with cool winters (mean 9.2°C), warm summers (mean 20.1°C), and annually variable precipitation that predominantly occurs in the winter and spring (averaging 622.5 mm per year) (Ackerly *et al*. 2002). The growing season occurs during the rainy season and plants senesce in early summer. This study was conducted during the growing seasons from mid-April to mid-May in 2015, 2016, and 2017. There was highly variable annual precipitation across the three years (491 mm, 604 mm, and 985 mm in 2015, 2016, and 2017, respectively; Weather Underground).

*S. pulchra* occurs at substantial densities in both serpentine and nonserpentine grasslands at JRBP (McNaughton 1968). The serpentine plant communities at JRBP are more diverse than the nearby nonserpentine plant communities and include many native grasses and forbs, in contrast to the invasive-dominated nonserpentine grasslands (McNaughton 1968; Huenneke *et al*. 1990; Field *et al*. 1996). Plant species commonly found in serpentine grasslands at JRBP include native *Stipa pulchra, Elymus glaucus, Elymus multisetus, Eschscholzia californica*, and the invasive grass *Bromus hordeaceus* (McNaughton 1968; Hobbs and Mooney 1985). The nonnative annual species that dominate the nonserpentine grasslands but are absent from serpentine grasslands include *Avena barbata, Avena fatua, Bromus diandrus*, and *Erodium botrys* (McNaughton 1968; Hobbs and Mooney 1985; Field *et al*. 1996). The serpentine and nonserpentine grasslands at JRBP share a land use history of livestock grazing that ceased in 1960, when conservation of these areas became a management priority (Bocek and Reese 1992).

All survey sites were located along the ridgetop at JRBP, where serpentine and nonserpentine (greenstone) soils are present in discrete adjacent bands (Fig. 1a) (Oze *et al*. 2004). Chemical analyses of the soils on this ridgetop show consistent, significant differences in serpentine and nonserpentine soil chemistry, with the serpentine soils enriched in heavy metals and trace elements (Fe, Cr, Ni, Mn, and Co), and impoverished in essential plant nutrients (Streit *et al*. 1993, Oze *et al*. 2004, 2008). The relative flatness of the ridgetop ensured that all survey sites were at roughly the same elevation and had similar slopes, aspects, and water availability (Oze *et al*. 2008).

**Figure 1.**
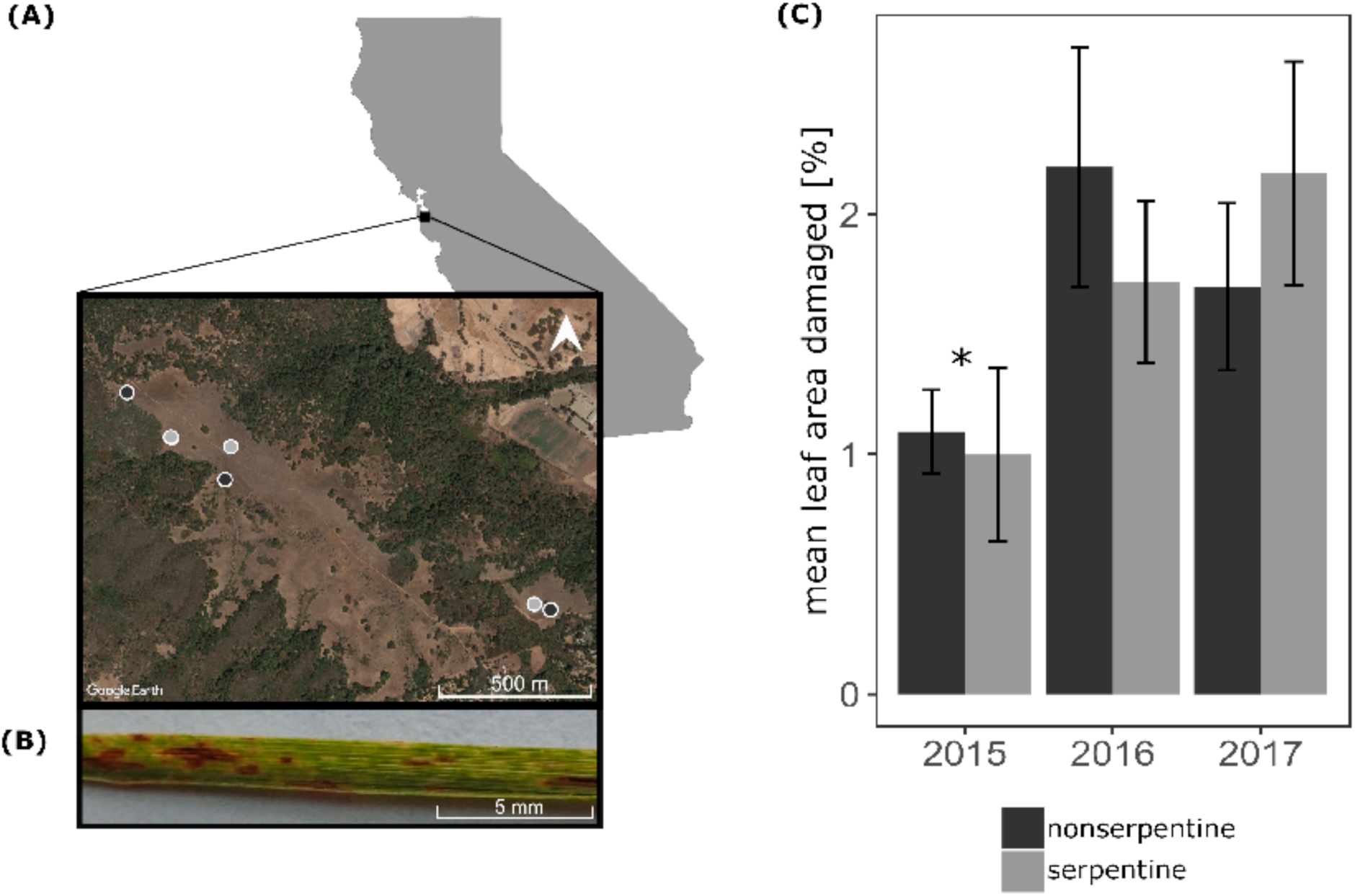
Pathogen damage on *Stipa pulchra* was consistently low across three years and both grassland types. **(A)** Locations of surveyed nonserpentine sites (blue) and serpentine sites (green) along the ridgetop at JRBP. **(B)** Example of fungal-caused pathogen foliar damage on *S. pulchra*. **(C)** Mean percentage (bars) and two standard errors (error bars) of diseased leaf area for surveyed serpentine and nonserpentine *S. pulchra*, by year. Damage was similar in both habitat types, but differed interannually. The asterisk indicates significantly lower damage in 2015 than in subsequent years of the survey.

### Quantification of disease

To assess plant disease burden, we quantified the percentage of living leaf area exhibiting symptoms of fungal disease in serpentine and nonserpentine populations of *S. pulchra*. We selected three serpentine grassland sites and three nonserpentine grassland sites such that each serpentine site was paired with a nearby (∼170m away) nonserpentine site (Fig. 1a). Individual sites within each habitat type were >300m apart and the maximum distance between any two sites was 1600m (Fig. 1a). Each year, we randomly placed four 5-meter transects at each 40m x 40m site and assessed the infection status of the *S. pulchra* individual nearest each meter mark, resulting in five individual plants per transect surveyed each year (N = 60 plants per habitat type per year). The infection status of individual plants was not tracked across years, although some plants could have been sampled repeatedly by chance. Following the protocol for estimation of foliar infection severity on grasses described in Mitchell *et al*. 2002 and Spear and Mordecai 2018, a single surveyor visually estimated the percentage of living foliar tissue damaged by pathogens for six arbitrarily-selected leaves per plant. We used printed field reference guides showing known amounts of damage on leaves to standardize estimates. Damage was indicated by spots and lesions which ranged in color from dark brown to pale yellow (Fig. 1b). Senesced tissue and herbivore damage (commonly in the form of scrapes and holes in the leaf surface) was excluded from the analyzed leaf area. For damage estimates of 5% or greater, values were determined to the nearest 5%. Damage levels below 5% were assigned values of 1% or 0.1%.

We used analysis of variance (ANOVA), with individual plants nested within sites and sites nested within habitat types, to test for separate effects of habitat type and year, as well as interactions between habitat type x year and habitat type x site, on log-transformed mean percent disease damage on serpentine and nonserpentine *S. pulchra* (N = 36 serpentine and 36 nonserpentine transects). When ANOVA results were statistically significant, we used t-tests for every possible pairwise combination, with Bonferroni adjustments of p-values to account for multiple comparisons, to identify the interactions contributing to the significant result.

### Isolation and molecular identification of foliar fungal pathogens

To identify the fungal foliar pathogens infecting serpentine and nonserpentine *S. pulchra* over all years, we harvested one segment of symptomatic leaf tissue for culturing and identification of the putatively causal fungal pathogen(s) from each individual included in the three-year damage survey with symptoms of disease. This tissue segment came from the first surveyed leaf with disease symptoms on each plant. We excised, surface sterilized, and plated a 4mm^2^ square of tissue from each sample at the leading edge of infection, to improve the chances of isolating the causal fungal pathogen, on malt extract agar with 2% chloramphenicol to prevent bacterial growth. While it is possible that the growing medium used may have led to selective isolation and sequencing of fungi other than those causing symptoms, our use of small segments of leaf tissue from the advancing margin of disease was intended to prevent this outcome under the assumption that symptom-causing organisms should be dominant in visibly diseased tissue and particularly at its expanding margin. We observed the plates for growth for eight weeks and isolated morphologically distinct hyphae into axenic culture. Six segments of surface sterilized, asymptomatic tissue were plated as a control and showed no growth. We extracted and sequenced genomic DNA from the ITS region for each isolate. Fungal isolation and sequencing methods with ITS1F and ITS4 primers followed Spear and Mordecai 2018.

We estimated taxonomic placement of fungal isolates by grouping sequences into operational taxonomic units with 97% sequence similarity (OTUs), a proxy for species delineation based on the range of intraspecific ITS divergence (O’Brien *et al*. 2005), and then comparing assembled DNA sequences to the entries in the UNITE fungal database. To prepare sequences for taxonomic placement, we first trimmed forward and reverse Sanger sequencing reads to a maximum error rate of 5% and then automatically assembled bidirectional reads into consensus sequences with a minimum of 20% overlap and 85% sequence similarity in Geneious (Kearse *et al*. 2012), discarding consensus sequences less than 450 bp in length and/or less than 60% in read quality. We grouped the remaining internal transcribed spacer (ITS) sequences into OTUs using USEARCH 10.0.240 (Edgar 2010) and estimated the taxonomic placement of each OTU in mothur v.1.40.5 (http://mothur.org/; Schloss 2009) using the naïve Bayes classification approach (classify.seqs with options: method = ‘wang’, cutoff = 0, iters = 200). We used a bootstrapping confidence score of at least 80% to assign a species name and at least 60% to assign taxonomy at higher ranks.

### Analyses of fungal community composition

To characterize foliar fungal community composition in serpentine and nonserpentine *S. pulchra* populations, we compared the diversity and assessed the similarity of communities of fungi cultured from the diseased tissue of surveyed plants. Additionally, we compared the identified fungal taxa to previously reported plant - fungal genus associations compiled in the USDA-ARS plant - fungal database (Farr and Rossman 2020), the Genera of Phytopathogenic Fungi (GOPHY) database (Marin-Felix *et al*., 2017, 2019), and the publications indexed in Web of Science and Google Scholar. To describe fungal community diversity, we calculated observed taxa richness and Fisher’s alpha, a widely used measure of species richness that is robust to uneven sample sizes (Fisher *et al*. 1943; Magurran 2013); generated taxa accumulation curves to understand sampling efficacy (Oksanen *et al*. 2016); and counted the number of fungal isolates in each genus, family, order, and class from each grassland type. We compared fungal community composition across grassland types and across years visually, using non-metric multidimensional scaling (NMDS), and statistically, using permutational multivariate analyses of variance (PERMANOVA) with pairwise Adonis functions for non-normalized OTU read counts in the R package vegan (Anderson 2001; Arbizu 2016; Oksanen *et al*. 2016). To assess spatial turnover in fungal communities, we included a site distance factor in the statistical analyses, with comparisons between sites <600m apart categorized as “near” and comparisons between sites >600m apart categorized as “far” (Fig. 1a). The PERMANOVA tested for effects of habitat type, year, distance between sites, and all possible interactions between each of these three factors, on fungal community composition. We used the function ‘betadisper’ with the ‘anova’ method to assess the potential for group differences in dispersion to affect the PERMANOVA results. To account for low culturing success in some transects, we considered the combined isolates from two nearby transects at each site in each year (N = 18 serpentine and 18 nonserpentine communities, generated from 36 total transects per habitat type) to be distinct communities for NMDS visualization.

For ANOVA and PERMANOVA analyses we considered the combined isolates from each site in each year to be distinct communities (N = 9 serpentine and 9 nonerspentine communities). When PERMANOVA results were statistically significant, we used pairwise PERMANOVAs with Bonferroni adjustments to p-values to clarify which fungal communities were significantly different from one another (Arbizu 2016). Additionally, we used the function ‘betadisper’ in vegan to assess the potential for differences in dispersion to affect the PERMANOVA results (Oksanen *et al*. 2016). For the NMDS and PERMANOVA analyses, we used the function vegdist with the abundance-based Chao method, accounting for unsampled species, to create a matrix of pairwise community dissimilarities (Chao *et al*. 2016; Oksanen *et al*. 2016). We also used the Morisita-Horn index, which is independent of sample size and diversity, to make pairwise comparisons of similarity within and between the serpentine and nonserpentine fungal communities from each year (N = 265 isolates, 200 bootstrap replicates), and within the entire fungal community between years (Wolda 1981; Jost *et al*. 2011). Finally, we used ANOVA to test for effects of habitat type, year, site distance, year x habitat type, and year x site distance interactions on fungal diversity (Fisher’s alpha) and richness at the site level.

To perform these fungal community analyses, we used the packages BiodiversityR (Kindt and Coe 2005), bipartite (Dormann *et al*. 2008), fossil (Vavrek 2011), rich (Rossi 2011), SpadeR (Chao *et al*. 2016), and vegan (Oksanen *et al*. 2016) with custom commands (Gardener 2014) in R version 3.4.1.

### Chemical analyses of leaf tissue

In 2016, we analyzed C and N content of the foliar tissue of the *S. pulchra* plants surveyed in serpentine and nonserpentine grasslands. Because both leaf age and infection status might influence leaf chemistry, we collected the youngest fully expanded and entirely asymptomatic leaf of each individual surveyed in the field (N = 60 serpentine, 58 nonserpentine). We dried the samples at room temperature for three months, then at 60°C for 48 hours, ground them to a powder, and then dried them again at 60°C for 24 hours. We measured C and N content of these samples with a Carlo Erba NA-1500 elemental analyzer using atropine standards.

To assess the potential for habitat type to mediate plant-pathogen interactions through its influence on plant tissue chemistry, we related habitat type to leaf tissue chemistry of *in situ S. pulchra*, and chemistry of these plants to disease burden and fungal species abundances. To relate leaf tissue chemistry to observed damage levels, we compared the mean percentage of C and N per dried weight of leaf tissue for surveyed serpentine and nonserpentine plants with Wilcoxon rank sum tests. We also compared mean C:N ratio, which is indicative of plant health (Ciompi *et al*. 1996; Brady *et al*. 2005). We used Pearson’s product-moment correlation to test for significant monotonic relationships between the log-transformed mean amount of diseased tissue observed on individual plants in the field and their foliar C and N content. Additionally, we used multivariate generalized linear models with resampling-based hypothesis testing to test for fungal community responses to tissue chemistry. For fungal species represented by five or more isolates in 2016, we tested for significant effects of foliar tissue C and N percentages, C:N ratio, and habitat type on fungal species observations. For these analyses, we used the functions ‘manyglm’ and ‘anova’ (nBoot = 999), with p-values adjusted for multiple comparisons, in the R package mvabund (Wang et al. 2012).

All statistical analyses were done in R version 3.4.1. In addition to the R packages specified above, we used the packages cowplot (Wilke 2019), data.table (Dowle *et al*. 2019), RColorBrewer (Neuwirth 2014), reshape2 (Wickham 2007), tidyverse (Wickham 2017), and xtable (Dahl *et al*. 2019) to manipulate, plot, and export data and results.

## Results

### Pathogen damage

Pathogen damage was similarly ubiquitous yet low-burden in serpentine and nonserpentine grassland habitats. Every plant surveyed exhibited evidence of foliar fungus-caused disease. The mean percentage of diseased leaf area observed across all years was 1.63 for serpentine and 1.66 for nonserpentine *S. pulchra*, respectively (Fig. 1c), and the mean proportion of surveyed leaves with pathogen damage was 0.83 for serpentine and 0.85 for nonserpentine plants. Year significantly affected log-transformed mean percent damage (ANOVA: df = 2, SS = 9.41, F-value = 5.375, p-value = 7.04 × 10^−3^) (Table 1). There were significant differences in percent damage between was between 2015 and 2016 and between 2015 and 2017 (2015 mean percent damage = 1.05; 2016 = 1.96; 2017 = 1.94; t-values = −2.66, −3.70; Bonferonni-adjusted p-values = 2.5 × 10^−3^, 8.1 × 10^−4^).

**Table 1.** Pathogen damage on *Stipa pulchra* varied significantly by year, but not by habitat type. ANOVA results for log-transformed transect-level measurements of percentage of symptomatic living tissue on serpentine and nonserpentine *S. pulchra* (N = 3 sites per habitat type per year, with 12 transects per site per year, for a total of 36 serpentine and 36 nonserpentine transects over all three years). The first column lists the factor(s) tested; the second column degrees of freedom (df); the third column the sum of squares (SS); the fourth column the mean squared error (MSE); the fifth column the F-value; and the 6^th^ column the p-value. P-values < 0.05 are marked with an asterisk.

| Variables | df | SS | MSE | F value | p-value |
| --- | --- | --- | --- | --- | --- |
| habitat type | 1 | 5.8 | 5.81 | 2.63 | 0.105 |
| year | 2 | 31.4 | 15.68 | 7.10 | 9.53 x 10<sup>-4</sup>* |
| habitat type x<br>site | 4 | 20.2 | 5.05 | 2.29 | 0.0598 |
| habitat type x<br>year | 2 | 9.9 | 4.96 | 2.25 | 0.107 |

### Fungal pathogen community

We isolated 267 unique fungal isolates from 258 plated symptomatic tissue pieces with fungal growth (144 nonserpentine and 114 serpentine tissue pieces from 36 nonserpentine and 34 serpentine transects, out of 360 total tissue pieces). Of the 267 unique fungal isolates, we successfully sequenced 256 isolates. The sequenced isolates clustered into 30 operational taxonomic units (OTUs) based on 97% sequence similarity, representing 23 genera, 15 families, 8 orders, and 5 fungal classes, primarily the Dothideomycetes. We were unable to identify 18 OTUs to the fungal species level, and four of these to the genus level, due to a lack of close sequence matches in the UNITE database and/or a lack of taxonomic information for close sequence matches (Table 2). The most common OTU was represented by 48 isolates and the least common OTUs were represented by one isolate each. OTUs are hereafter referred to as “species” (O’Brien et al. 2005).

**Table 2.** Fungal species isolated from symptomatic *Stipa pulchra* leaves were unevenly distributed across habitat types. Each row represents one of the species isolated. The columns indicate the isolation frequency for each species in each soil type. Asterisks indicate fungal species in genera not previously reported from plant hosts in the Poaceae; daggers indicate species not previously reported in association with any plant disease symptoms; double daggers indicate species in genera not previously reported in association with foliar disease symptoms.

| **Species** | **Nonserpentine** | **Serpentine** |
| --- | --- | --- |
| *Alternaria* cf. *infectoria* | 26 | 21 |
| *Keissleriella* sp. † | 19 | 8 |
| *Drechslera* cf. *fugax* | 17 | 1 |
| *Stemphylium* sp. | 15 | 8 |
| *Ramularia* cf. *proteae* | 11 | 37 |
| *Drechslera* sp. | 9 | 1 |
| *Neoascochyta* sp. | 7 | 3 |
| *Cladosporium* sp. | 6 | 3 |
| *Alternaria* cf. *eureka* | 5 | 2 |
| *Parastagonospora* cf. *avenae* | 5 | 0 |
| *Pyrenophora* cf. *chaetomioides* | 5 | 2 |
| *Pyrenophora* cf. *lolii* | 4 | 3 |
| *Phaeosphaeriaceae* sp. 2 | 3 | 5 |
| *Pleosporaceae* sp. | 3 | 2 |
| *Drechslera* cf. *nobleae* | 1 | 5 |
| *Epicoccum* sp. | 1 | 0 |
| *Heyderia* sp. *† | 1 | 1 |
| *Parastagonospora* cf. *cumpignensis* | 1 | 0 |
| *Parastagonospora* sp. | 1 | 0 |
| *Phaeosphaeria* sp. | 1 | 2 |
| *Podospora* sp. † | 1 | 0 |
| *Phaeosphaeriaceae* sp. 1 | 1 | 0 |
| *Chalastospora* cf. *gossypii* ‡ | 0 | 1 |
| *Hyphodiscus* cf. *brevicollaris* *† | 0 | 1 |
| *Nemania* cf. *serpens* *‡ | 0 | 1 |
| *Phaeophlebiopsis* sp. *† | 0 | 1 |
| *Plectania* sp. *† | 0 | 2 |
| *Pseudoseptoria* sp. | 0 | 1 |
| *Helotiales* sp. | 0 | 1 |
| *Wojnowiciella* sp. * | 0 | 1 |

All genera identified include fungal species previously isolated from plant hosts, and the majority of these genera include known plant pathogens (Table 2). The taxonomic results indicate that many of the species represent close relatives of major agricultural plant pathogens including *Alternaria, Cladosporium, Drechslera, Phaeosphaeria, Pyrenophora, Stemphylium, Ramularia*, and *Wojnowiciella* spp. (Table 2) (Kirk *et al*. 2008; Stukenbrock and McDonald 2008; Havis *et al*. 2015; Marin-Felix *et al*. 2017, 2019; Farr and Rossman 2020). Inoculation experiments described in Spear and Mordecai 2018 previously confirmed the pathogenicity of *Parastagonospora* and *Pyrenophera* species on *S. pulchra* leaves. Six genera, *Heyderia, Hyphodiscus, Nemania, Phaeophlebiopsis, Plectania*, and *Wojnowiciella*, are here reported for the first time from hosts in the Poaceae; six genera, *Heyderia, Hyphodiscus, Keissleriella, Phaeophlebiopsis, Plectania*, and *Podospora* are reported for the first time in association with plant disease symptoms; and two additional genera, *Chalastospora* and *Nemania*, are reported for the first time in association with foliar disease symptoms (Table 2) (United States Department of Agriculture 1960; Goetz and Dugan 2006; Marin-Felix *et al*. 2017, 2019; Golzar *et al*. 2019; Adikaram and Yakandawala 2020; Farr and Rossman 2020).

Fungal community diversity was similar between serpentine and nonserpentine habitats and between years (Table 2). Although fungal richness was higher at every taxonomic level in the serpentine relative to the nonserpentine, these differences were not statistically significant (Tables 2, S1-S5). Twenty-four fungal species were isolated from serpentine *S. pulchra* (Fisher’s alpha = 9.324, 95% CI = 4.664, 13.983) and 22 from nonserpentine (Fisher’s alpha = 7.261, 95% CI = 3.597, 10.925); 16 of these species (53%) were shared (Fig. 2; Tables 1, 2). Habitat type, year, and year x habitat type were not significantly related to fungal richness or Fisher’s alpha at the site level (ANOVA: Tables S6, S7). Novel genera were isolated for both community types in every year of surveying, with only seven of 30 (23%) observed species and six of 23 genera (26%) isolated in all three years (Table S5). Species accumulation curves did not approach horizontal asymptotes for either habitat type, suggesting that neither fungal community was fully described (Fig. S1).

**Figure 2.**
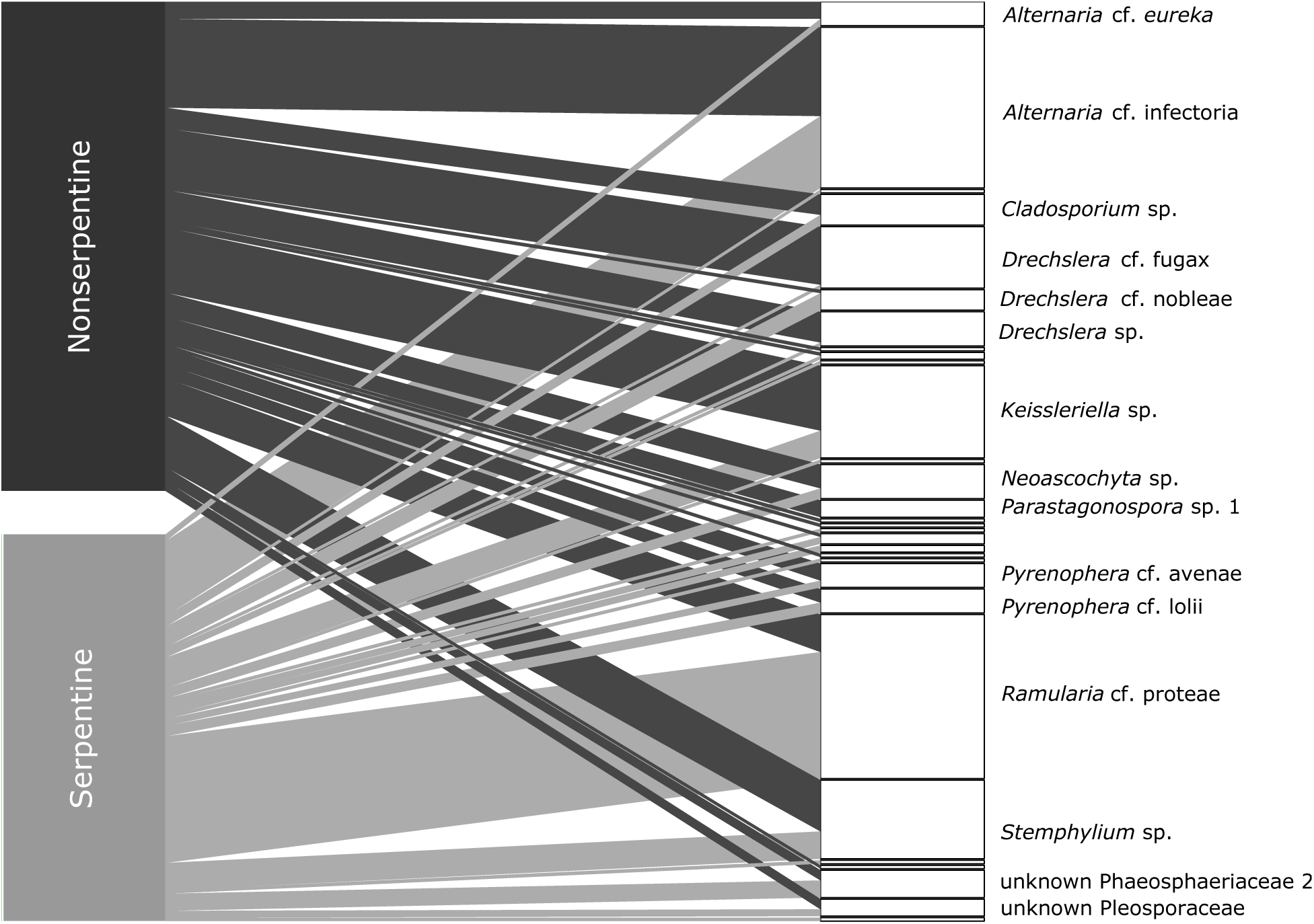
Pathogens isolated from *Stipa pulchra* growing in nonserpentine and serpentine grasslands. Bipartite network showing species interactions between *S. pulchra* plants (left) growing in serpentine grasslands (green) and nonserpentine grasslands (blue) and fungal species cultured from symptomatic leaf tissue (white bars on the right). The length of each white bar indicates the number of isolates in the operational species unit (based on 97% sequence similarity). The thickness of lines connecting left and right bars represents the number of times a particular fungal species was isolated from *S. pulchra* in a particular grassland type. The nonserpentine fungi came from 15 genera, 8 families, 4 orders, and 3 classes; the serpentine fungi came from 20 genera, 14 families, 7 orders, and 5 classes (Tables 1, S1-S5).

Both fungal communities were dominated by a few abundant species (isolated >10 times), but the most abundant species differed between nonserpentine and serpentine habitats: these were *Alternaria* cf. *infectoria* and *Ramularia* cf. *proteae*, respectively (Table 2; Fig. S2). The nonserpentine community included five abundant species, representing 62% of the isolates, while the serpentine community included two abundant species, representing 51% of serpentine isolates (Table 2; Fig S2). The serpentine community had a higher proportion of rare species (<3 isolates) than the nonserpentine community, 63% versus 36%, respectively. One *Drechslera* species was abundant in the nonserpentine community and rare in the serpentine (Table 2; Fig. S2).

Habitat type and year significantly affected fungal community composition, based on two-way PERMANOVA analysis of site-level fungal communities and NMDS visualization of community similarity (habitat type: F-value = 5.112, R^2^ = 0.163, p = 0.003; year: F-value = 5.655, R^2^ = 0.360, p = 0.001; Fig. 3; Table 4). Fungal communities had homogeneous dispersions across habitat types (average distance to median: serpentine = 0.53, nonserpentine = 0.56; ANOVA F-value = 1.10, p-value = 0.3) and years (average distance to median: 2015 = 0.38, 2016 = 0.34, 2017 = 0.27; ANOVA F-value = 0.93, p-value = 0.42), indicating that significant PERMANOVA results were due to true differences in centroid positions. Serpentine fungal communities were generally more similar to one another than they were to nonserpentine fungal communities, and vice versa (Fig. 3; Tables 2, 3). Community composition differed significantly between 2015 and 2017, and 2016 and 2017, but not between 2015 and 2016 (Table S8). Differences in fungal community composition between years did not correspond to differences in disease severity between years (Figure 1c, Table S8).

**Table 3.** Fungal communities were distinct on *Stipa pulchra* plants growing in serpentine versus nonserpentine grasslands, across all years. Fungal community similarity by habitat type and year. The leftmost column lists the fungal communities considered, with the number of species in each community in parentheses. The second column lists Fisher’s alpha for each community. The third column shows the number of shared species and, in parentheses, the percentage of total observed species that were shared for each pair of communities. The rightmost column lists the estimated Morisita-Horn community overlap value, based on absolute species abundances, for each pair of communities. The Morisita-Horn index ranges from 0 to 1, with 1 indicating complete overlap.

| **Fungal community pair** | **Fisher's alpha** | **Shared species** | **Morisita-Horn Similarity Index** |
| --- | --- | --- | --- |
| All years serpentine (24) | 9.324 | 16 | 0.675 |
| All years nonserpentine (22) | 7.261 | (53.0%) |  |
| 2015 serpentine (11) | 7.955 | 5 | 0.535 |
| 2015 nonserpentine (12) | 5.205 | (27.8%) |  |
| 2016 serpentine (16) | 7.897 | 11 | 0.766 |
| 2016 nonserpentine (13) | 5.861 | (61.1%) |  |
| 2017 serpentine (12) | 6.784 | 8 | 0.725 |
| 2017 nonserpentine (12) | 5.136 | (50.0%) |  |
| 2015 serpentine (11) | 7.955 | 7 | 0.707 |
| 2016 serpentine (16) | 7.897 | (35.0%) |  |
| 2015 serpentine (11) | 7.955 | 5 | 0.721 |
| 2017 serpentine (12) | 6.784 | (27.8%) |  |
| 2016 serpentine (16) | 7.897 | 7 | 0.786 |
| 2017 serpentine (12) | 6.784 | (33.3%) |  |
| 2015 nonserpentine (12) | 5.205 | 6 | 0.668 |
| 2016 nonserpentine (13) | 5.861 | (31.5%) |  |
| 2015 nonserpentine (12) | 5.205 | 5 | 0.295 |
| 2017 nonserpentine (12) | 5.136 | (26.3%) |  |
| 2016 nonserpentine (13) | 5.861 | 7 | 0.882 |
| 2017 nonserpentine (12) | 5.136 | (38.9%) |  |

**Table 4.**
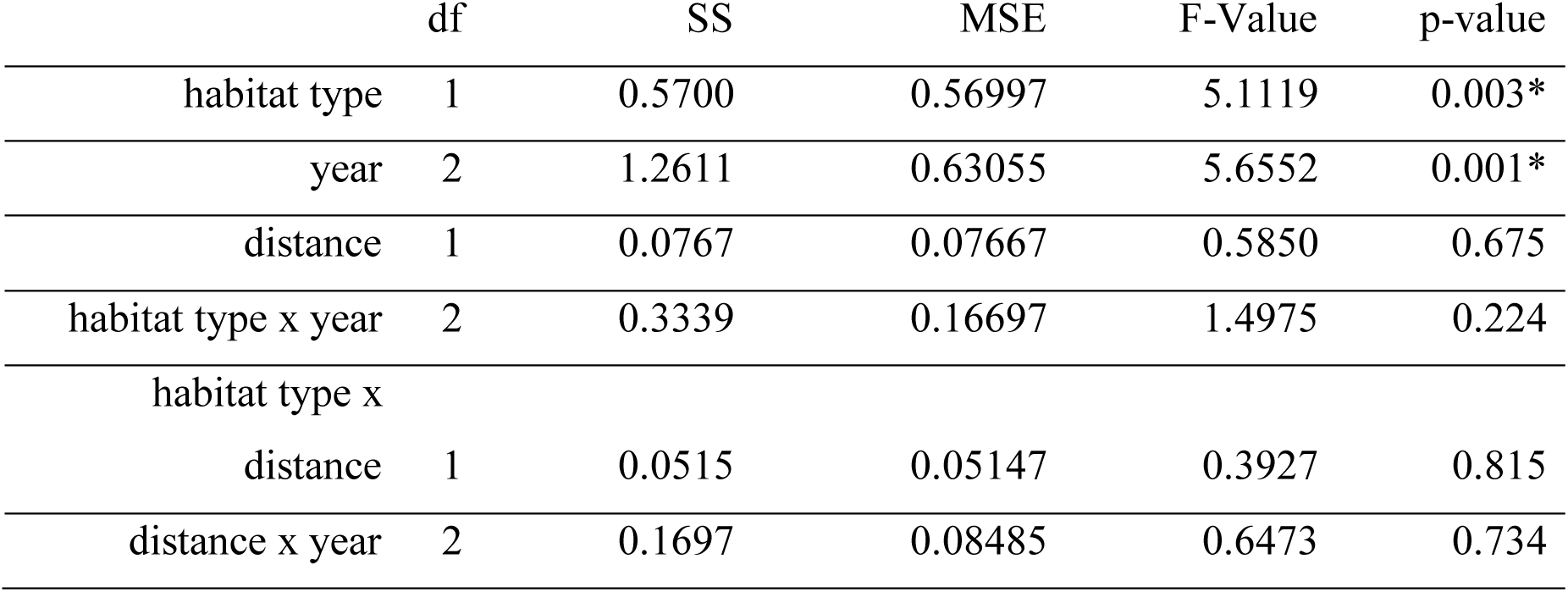
PERMANOVA results show significant effects of both habitat type and year on fungal foliar community composition (N = 18 serpentine and 18 nonserpentine sites). From left to right, columns list factors tested, degrees of freedom (df), sum of squares (SS), mean squared error (MSE), F-value, and p-value. Statistically significant p-values (p < 0.05) are marked with an asterisk. The distance factor compares communities at near (<600m) vs. far (>600m) distances from one another.

**Figure 3.**
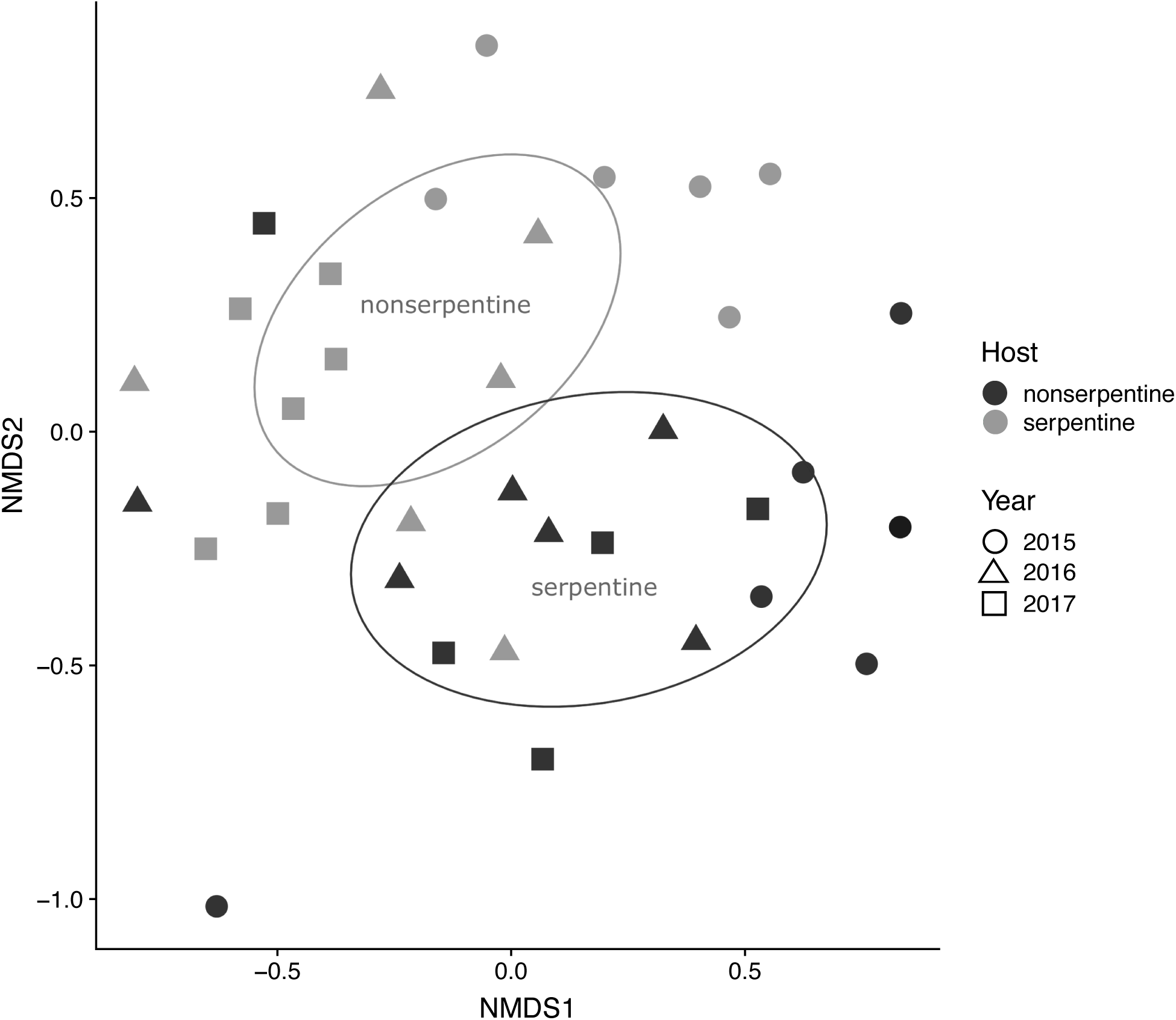
Distinct fungal communities were isolated from *Stipa pulchra* growing in nearby serpentine and nonserpentine sites across years. Non-metric multidimensional scaling visualization of serpentine and nonserpentine fungal community similarity. Each point corresponds to the combined fungal community of two transects at the same site in the same year. Serpentine communities are shown in green and nonserpentine communities are shown in blue. Circles, triangles, and squares represent fungal communities sampled in 2015, 2016, and 2017, respectively. Ellipses enclose 95% confidence intervals for ordination of serpentine and nonserpentine communities. Community similarity decreases with the distance between points. PERMANOVA analysis showed a significant effect of soil type on fungal community composition (F = 3.623, R^2^ = 0.0618, p = 0.001).

Interpretation of the statistical significance of differences in Morisita-Horn community overlap that fall between zero (no overlap) and one (complete overlap) is a longstanding challenge in community ecology (Horn 1966, Rodrigues and Vieira 2010). Qualitatively, estimates of the Morisita-Horn similarity index for communities within habitat types suggest that the amount of interannual species turnover is more consistent within serpentine than nonserpentine communities (Fig. 3, Tables 2, 3). While serpentine community overlap between years ranged from 0.707 to 0.786, nonserpentine community overlap ranged between 0.295 and 0.882 (Table 3). Within-year variation in species turnover between habitat types was comparatively moderate, with Morisita-Horn overlap ranging from 0.535 to 0.725 (Table 3).

### Leaf tissue chemistry and plant – pathogen interactions

As expected from differences in soil type between habitats, leaf chemistry in plants occurring on serpentine versus nonserpentine soils differed in mean C and N content. Serpentine plants had significantly lower mean C and N content than nonserpentine plants (Fig. 4; Table S9). C:N ratio was similar for plants in both habitat types. Log-transformed mean percent symptomatic tissue was not significantly associated with percent C, percent N, or C:N ratio in asymptomatic foliar tissue for serpentine or nonserpentine plants, based on Pearson’s product-moment correlation tests (Fig. 4). For the fungal community consisting of the seven species isolated five or more times in 2016 (N = 74 isolates), all of which occurred at least once in each habitat type, species composition differed between habitats, but was not associated with foliar percent C, percent N, or C:N ratio (Table S10).

**Figure 4.**
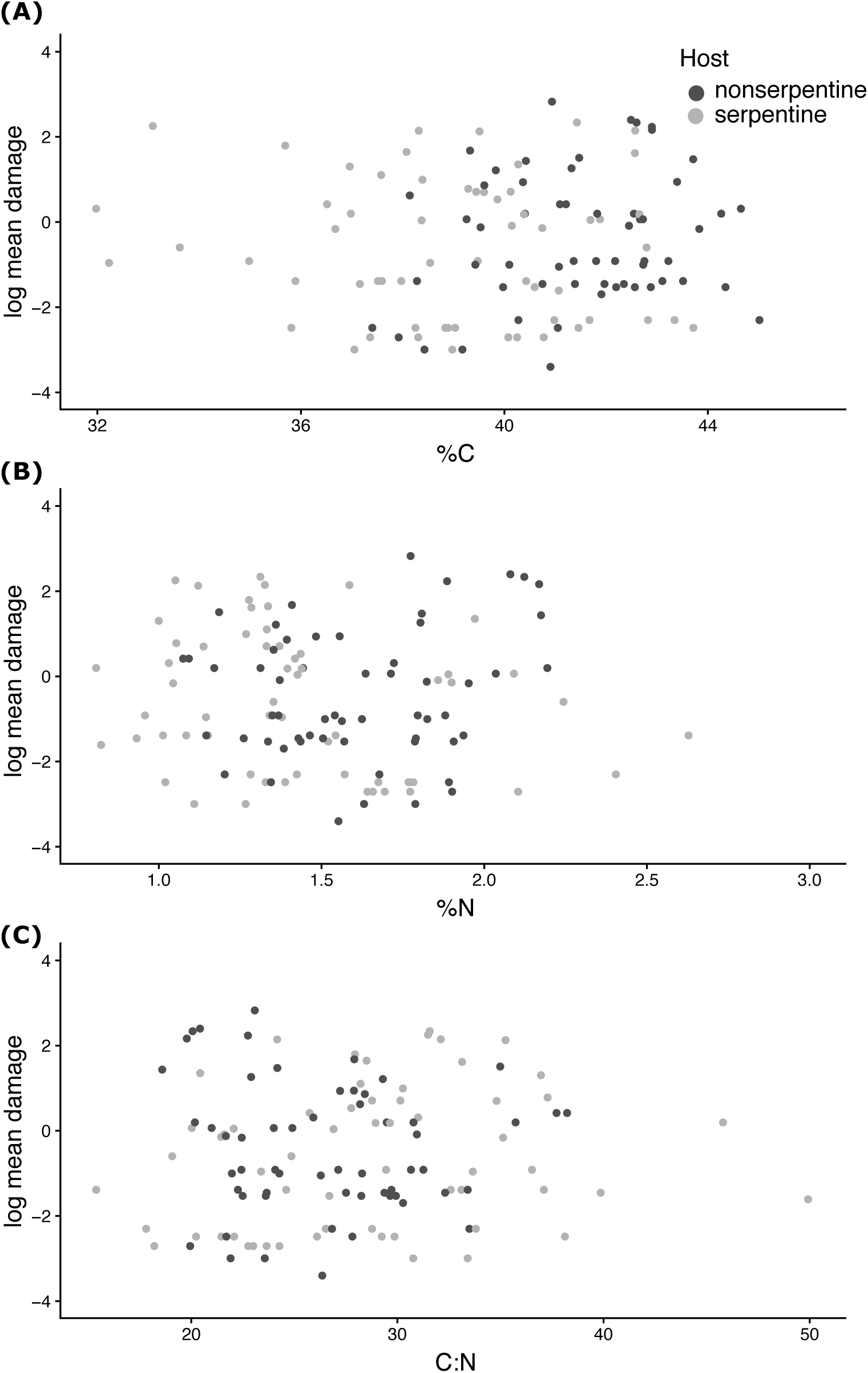
*Stipa pulchra* foliar C and N content differs between habitat types, but does not predict mean pathogen damage. **(A)** Foliar percent C, **(B)** percent N, and **(C)** C:N ratio versus mean percentage of diseased leaf area for serpentine and nonserpentine *S. pulchra* plants. Wilcoxon rank-sum tests indicated significantly higher foliar percent C and foliar percent N in nonserpentine compared to serpentine plants. Pearson’s correlation tests showed no relationships between foliar tissue chemistry and the amount of symptomatic leaf tissue observed on individual plants.

## Discussion

Here we show that a diverse suite of foliar fungi is associated with ubiquitous, low-burden damage on host individuals across growing seasons and habitat types. Habitat type appeared to shape fungal community composition in this system, but not foliar disease burden. In contrast, sampling year affected both fungal community composition and disease burden. However, this interannual variation in disease burden was not mediated by differences in pathogen community composition (Figs. 1c, 3; Table S8). While C and N content measured in young, healthy leaves differed between habitat types, variation in leaf chemistry did not explain variation in infection severity or in fungal community composition (Fig. 4; Tables S9-10). The results provide some of the first evidence that local growing conditions structure the pathogen communities infecting a single host species even over relatively small areas of a few hundred meters, where hosts and pathogens experience similar climate conditions (Figs. 1a, 3; Table 3). Overall, the findings imply that environmental heterogeneity within landscapes may both generate barriers to fungal disease spread in wild plant populations and bolster fungal diversity.

### Foliar disease burden

Mean levels of foliar infection were low even at their peak of 1.9% symptomatic leaf area in 2017 (Fig. 1c; Table 1). This level of damage is similar to that observed on other grass species in nonserpentine grasslands at JRBP (Spear and Mordecai 2018), and on taxonomically diverse tropical forest plants in Mexico and Hawaii (García-Guzmán and Dirzo 2001, Funk and Throop 2010). Recent work in the JRBP grassland system shows that symptoms in the amounts observed inflict only minimal plant fitness costs, suggesting that *S. pulchra* is able to effectively resist or suppress damage by foliar pathogens in both serpentine and nonserpentine grasslands (Spear and Mordecai 2018; Uricchio *et al*. 2019). These expectations align with assessments in wheat – fungal pathogen systems that indicate yield reductions of ≤ 1% for each percent of symptomatic leaf area (Milus 1993, Khan *et al*. 1997). The finding that between-habitat differences in foliar C and N content do not correspond to differences in infection severity aligns with the finding that infection severity did not differ between habitats (Figs. 1, 4). Additionally, similar C:N ratios suggesting that serpentine and nonserpentine plants are equally healthy correspond to the consistently low damage observed across habitats (Fig. 4; Table S9). However, the observational study design and use of young, healthy leaves for the tissue chemistry measurements limit interpretation of the observed relationships between leaf chemistry and disease severity. Because both tissue age and fungal infections can affect plant chemistry (Paul 1989; Donaldson *et al*. 2006), the leaf chemistry measurements may not accurately represent the chemical environment encountered by the pathogens that caused the observed damage.

### Fungal community composition

Our results suggest that habitat filtering, rather than dispersal limitation, plays a key role in fungal community assembly. The sampled nonserpentine and serpentine plants are in close proximity, yet habitat type—not distance between sites—predicted fungal community composition (Fig. 3; Tables 4, S10) (Martiny *et al*. 2006). Fungal community composition, and its relative stability over time, varied between habitats (Figs. 1c, 2, 3, S2; Tables 1-4). The results imply that abiotic and biotic differences in local environmental conditions can lead to differences in plant-fungal interactions in distinct habitat types. We hypothesize that influential habitat characteristics in this system include differences in pathogen sharing among the predominantly native hosts in serpentine grasslands versus the predominantly nonnative hosts in nonserpentine grasslands, in plant phenotypes, and/or in plant genotypes.

Plant hosts may differ in competence for different fungal species, and fungal species may in turn vary in transmissibility to *S. pulchra*. These two factors, and their interaction, likely contribute to differences in pathogen community composition on *S. pulchra* between serpentine and nonserpentine habitats. Serpentine grassland plant communities are generally more taxonomically diverse and stable across years than nonserpentine grasslands (Fernandez-Going *et al*. 2012). Qualitatively, our results suggest that fungal communities follow the same pattern of higher diversity and lower species turnover between years in serpentine areas, indicating potential links between plant and pathogen community structures (Fig. S2; Tables 1, 3, S1-S5). Research on five common invasive and native grasses (including *S. pulchra*) at JRBP shows that multi-host foliar fungal pathogens dominate this system, and that *S. pulchra* shares different pathogens with different co-occurring plant species, supporting this hypothesis (Spear and Mordecai 2018).

Differing nonserpentine and serpentine *S. pulchra* phenotypes might also influence fungal community assembly through a variety of mechanisms, including impacts of plant chemistry on plant immune pathways and/or fungal pathogen virulence, growth, and reproduction. Although differences in plant C and N content did not affect the distributions of commonly isolated fungal species found in both habitat types (Tables 2, S10), the limitations of the foliar chemistry measurements described above apply equally here. Importantly, the impacts of leaf C and N content on the distributions of rare species, many of which appear to be habitat specialists, remain unassessed (Tables 2, S10). Evidence that serpentine soil chemistry impacts on plant Ca content affect the frequency and severity of *Hesperolinon spp*. infection by the fungal rust pathogen *Melampsora lini* indicates the possible importance of habitat influence on elements other than C and N in *S. pulchra* tissues (Springer *et al*. 2006; Springer 2009). Potted *S. pulchra* plants grown from seed and serpentine soils collected in these sites had significantly lower Ca and P content and higher Ni and Mg content than similar plants grown in greenhouse soils, potentially suggesting additional phenotypic differences between serpentine and nonserpentine plants that may affect fungal community composition (Farner *et al*., unpublished data).

Overall, further investigation is needed to test the hypothesis that habitat-associated plant phenotypic differences shape *S. pulchra* foliar fungal communities. Future studies should include analyses of more fungal species, more elements, and more plant traits, including leaf physiological characteristics that might respond to elevated water stress in serpentine habitats. Based on the genetic foundations of the complex set of physical, chemical, and genetic defenses that define plant immune profiles, investigation into the extent of genetic divergence between serpentine and nonserpentine *S. pulchra* could also clarify the mechanisms that structure its fungal communities (Dodds and Rathjen 2010; Łaźniewska *et al*. 2012). While there are no physical barriers to prevent gene flow between individuals in different habitat types, the occurrence of phylogenetically and chemically distinct serpentine and nonserpentine populations of the native forb *Lasthenia californica* occurring in the same grasslands at JRBP suggests the potential for similar structure in the focal *S. pulchra* population (Desrochers and Bohm 1993; Chan *et al*. 2002; Rajakaruna *et al*. 2003).

In addition to turnover between habitat types, interannual fungal species turnover contributed substantially to fungal diversity in this system, with only seven of 30 total species isolated in all three years (Fig. 3; Tables 1, S5). Interannual variation in these communities may be driven by a variety of mechanisms, including high yearly variation in precipitation (2015 rainfall = 491 mm; 2016 = 604 mm, 2017 = 985 mm) (Weather Underground), plant population sizes, occurrences of different plant genotypes, and plant community composition (Hobbs and Mooney 1991). Although few studies track interannual variation in fungal communities on plants, the year-to-year dissimilarity of those hosted by *S. pulchra* is similar to that observed across a diverse taxonomic range of plants including ferns, legumes, and shrubs (Del Olmo-Ruiz and Arnold 2014; Cotton *et al*. 2015; Zhang *et al*. 2016). In this context, our results are consistent with a growing body of evidence for the general importance of storage effects—in which temporally variable conditions act as ecological niches that facilitate species diversity—in plant-associated fungal communities (Warner and Chesson 1985). This finding highlights the necessity of long-term data collection to accurately characterize plant-fungal interaction networks.

The variation in fungal community composition we observed on a single host species with habitat type and over time suggests possible mechanisms preventing severe outbreaks of foliar disease in California grasslands, where uniformly low disease burden was observed in this, and other, studies (e.g., Spear and Mordecai 2018). Transitions between distinct soil types over distances within the range of fungal spore dispersal may prevent the spread of pathogens that are sensitive to leaf tissue chemistry or have limited host ranges. Interannually variable conditions favoring successful infection by different fungal species each growing season suggest temporal barriers to pathogens evolving high virulence on both nonserpentine and serpentine plants (Table S5). Clarifying the specific mechanisms that drive differences in serpentine and nonserpentine fungal communities will require inoculation experiments to test how plant susceptibility to infection by different fungal species changes with plant tissue chemistry and plant genetics. Characterization of comprehensive plant-pathogen interaction networks including all plant species co-occurring with *S. pulchra* in serpentine and nonserpentine grasslands will improve our understanding of the role of the surrounding plant community in structuring fungal pathogen communities.

### Fungal diversity

Our work underscores the sparsity of current scientific understanding of plant – fungal interactions in natural systems. The novel plant host and plant disease associations reported here for nine fungal genera highlight the wealth of plant – fungal interactions that remain to be described (Table 2). Because this study is observational, the pathogenicity on *S. pulchra* of many of the fungal species isolated—even those from genera including well-known plant pathogens— remains unknown. It is possible that some of the isolated fungal taxa are not primary pathogens on this species, but endophytes that opportunistically act as secondary pathogens. Inoculation experiments and high-throughput sequencing studies to characterize and compare entire fungal communities inhabiting symptomatic and healthy *S. pulchra* leaf tissue will shed light on the ecology of the isolated taxa and on the potential for interactions between co-occurring fungal species to impact plant health.

These open questions notwithstanding, *S. pulchra* appears to host an unusually species-rich community of culturable fungi within its diseased foliar tissue when compared to other plant species’ interactions with both pathogenic and non-pathogenic fungi. Where we isolated 30 fungal species, past studies that described fungal pathogen diversity for more than 100 different plant hosts consistently found fewer than 20 fungal pathogens per species, and almost all hosts had fewer than ten pathogens (Mitchell and Power 2003; Hantsch *et al*. 2013; Rottstock *et al*. 2014). The level of fungal diversity we documented is also high compared to that of culturable fungal endophyte communities associated with asymptomatic plant tissues: of 35 plant species analyzed by Arnold and colleagues, only five were found to host 30 or more endopyhtes (Arnold and Lutzoni 2007; Del Olmo-Ruiz and Arnold 2014). Notably, studies that utilize high-throughput sequencing techniques to observe both culturable and unculturable endophytic fungi frequently report hundreds of fungal OTUs colonizing single host species (e.g., Jumpponen and Jones 2009, Zimmerman and Vitousek 2012). However, such results are difficult to compare to our own given the mismatch in the detection capabilities of the methods used. Nonetheless, the exceptional diversity of the culturable foliar fungi that inhabit *S. pulchra* is further indicated by species accumulation curves that suggest many more species in this community remain to be discovered (Fig. S1), by the lack of close sequence matches in genetic databases for some of the species isolated (Table 2), and by the isolation of multiple genera previously unassociated with plant hosts in the Poaceae (Table 2). These findings suggest that the characteristics that make the California Floristic Province a biodiversity hotspot may extend to the fungal diversity associated with this flora (Myers *et al*. 2000).

### Conclusions

Fungi associated with foliar disease in the native bunchgrass *S. pulchra* are relatively benign, likely cause minimal fitness impacts (Spear and Mordecai 2018), and contribute substantially to the overall biodiversity of California grasslands. Variation between habitats appears to play an important role in supporting the fungal diversity observed here, and may also reduce disease risk for *S. pulchra*. The high taxonomic diversity and rarity of serpentine fungi demonstrates that this community is potentially at risk of species loss via host-parasite coextinction as nonnative annual grasses continue to expand their ranges at the cost of native plants, and human activities including development and increased C and N deposition contribute to the loss of serpentine grasslands that already make up <2% of California’s surface area (Huenneke *et al*. 1990; Vallano *et al*. 2012). This work demonstrates that, in addition to >200 endemic plant species, California’s serpentine grasslands support unique communities of leaf-infecting fungi, providing additional motivation for active conservation of native grassland communities based on the far-reaching loss of biodiversity across trophic levels associated with their disappearance (Sprent 1992; Dunn *et al*. 2009).

## Acknowledgements

We thank Nona Chiariello, Caroline Daws, Joe Wan, Peter Vitousek, Scott Fendorf, Lawrence Uricchio, Guangchao Li, Douglas Turner, Virginia Walbot, Cody Hamilton, and the Mordecai and Peay Lab Groups for their help. This work was supported by the Jasper Ridge Kennedy Endowment, the National Science Foundation (DEB-1518681), Stanford Vice Provost for Undergraduate Education, and Stanford Bio-X. JEF and EAM were supported by the National Institutes of Health (R35GM133439).

## Supporting Information

**Figure S1.**
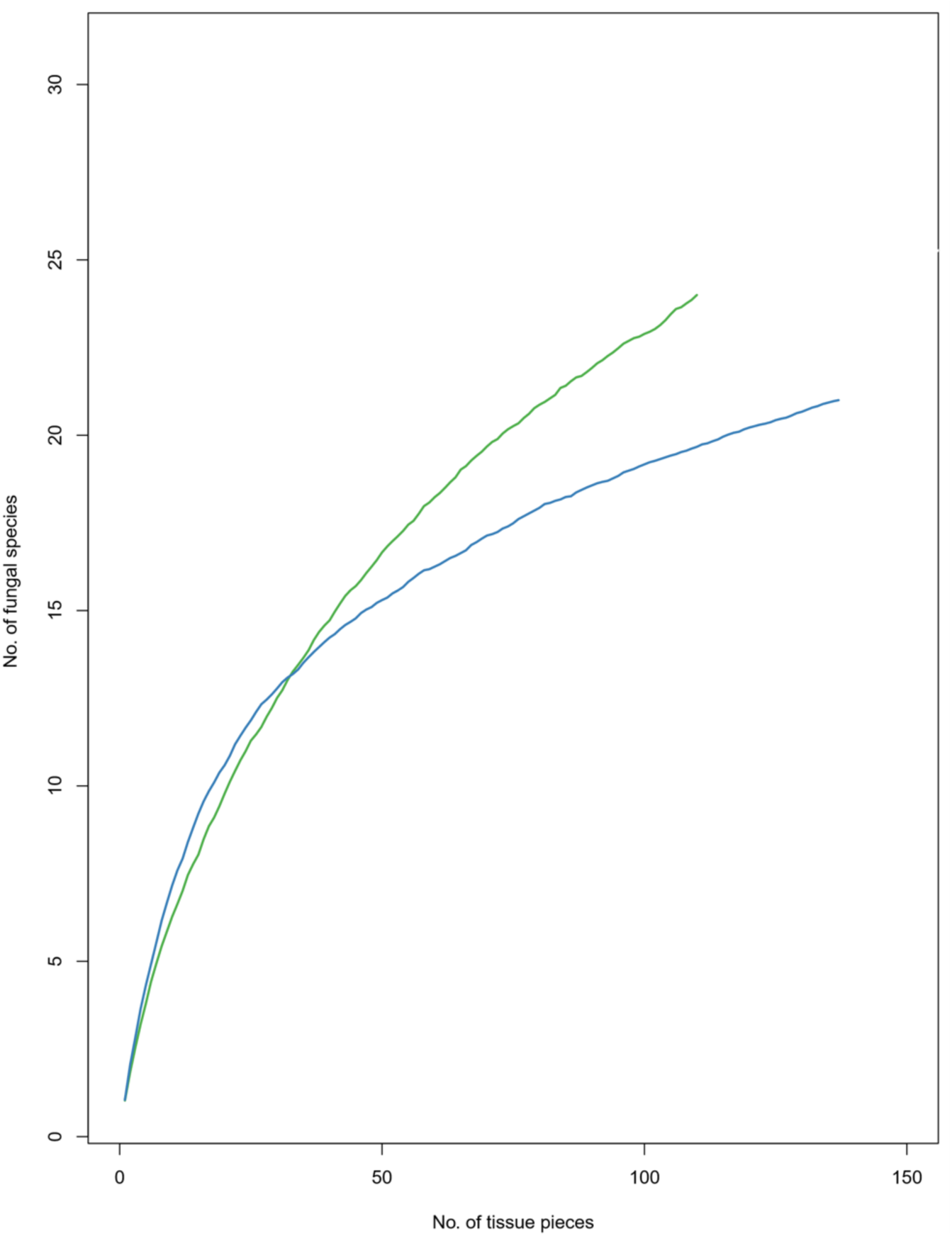
Species accumulation curves for serpentine (green) and nonserpentine (blue) fungal communities isolated from *Stipa pulchra* leaves over three years of surveys. The cumulative number of unique fungal OTUs (based on 97% sequence similarity; y-axis) is plotted against the number of *S. pulchra* leaf tissue pieces sampled (x-axis).

**Figure S2.**
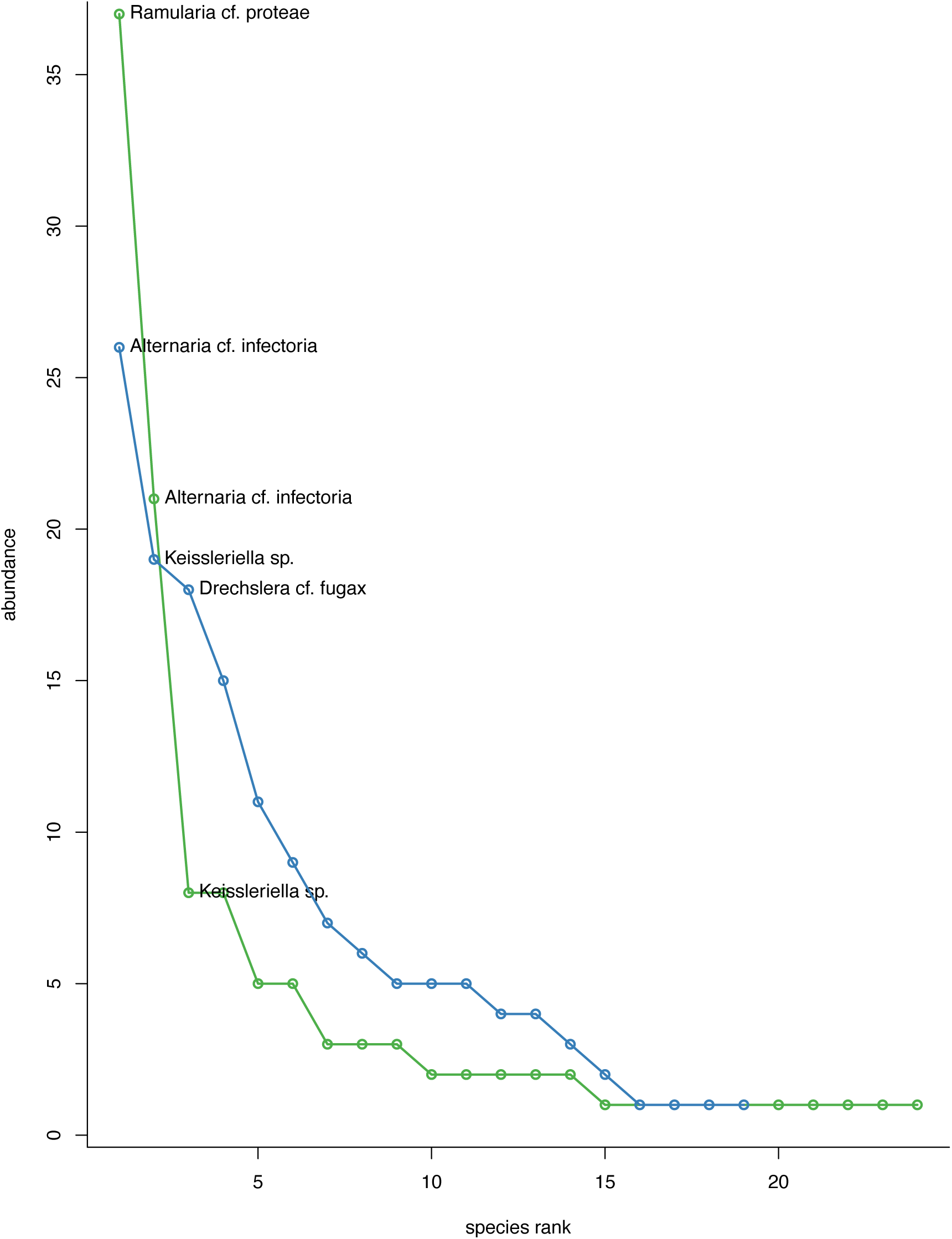
Fungal species abundance by rank for serpentine (green) and nonserpentine (blue) fungal communities on *Stipa pulchra* leaves. Each point represents the number of times a single species was isolated from *S. pulchra* leaves in each habitat type.

**Table S1.** Genus counts for serpentine and nonserpentine fungal communities isolated from *Stipa pulchra* leaves. Numbers in each column represent the number of times a genus was isolated from *S. pulchra* leaves in each habitat type.

| **Genus** | **Nonserpentine** | **Serpentine** |
| --- | --- | --- |
| *Alternaria* | 31 | 23 |
| *Drechslera* | 27 | 7 |
| *Keissleriella* | 19 | 8 |
| *Stemphylium* | 15 | 8 |
| *Ramularia* | 11 | 37 |
| *Pyrenophora* | 9 | 5 |
| *Neosascochyta* | 7 | 3 |
| *Parastagonospora* | 7 | 0 |
| *Cladosporium* | 6 | 3 |
| Phaeosphaeriaceae | 4 | 5 |
| unknown Pleosporaceae | 3 | 2 |
| *Epicoccum* | 1 | 0 |
| *Heyderia* | 1 | 1 |
| *Phaeosphaeria* | 1 | 2 |
| *Podospora* | 1 | 0 |
| *Chalastospora* | 0 | 1 |
| *Hyphodiscus* | 0 | 1 |
| *Nemania* | 0 | 1 |
| *Phaeophlebiopsis* | 0 | 1 |
| *Plectania* | 0 | 2 |
| *Pseudoseptoria* | 0 | 1 |
| unkown Helotiales | 0 | 1 |
| *Wojnowiciella* | 0 | 1 |

**Table S2.** Family counts for nonserpentine and serpentine fungal communities isolated from *Stipa pulchra* leaves. Numbers in each column represent the number of times a family was isolated from *S. pulchra* leaves in each habitat type.

| **Family** | **Nonserpentine** | **Serpentine** |
| --- | --- | --- |
| Pleosporaceae | 85 | 45 |
| Lentitheciaceae | 19 | 8 |
| Phaeosphaeriaceae | 12 | 8 |
| Mycosphaerellaceae | 11 | 37 |
| Didymellaceae | 8 | 3 |
| Cladosporiaceae | 6 | 3 |
| Helotiaceae | 1 | 1 |
| Lasiosphaeriaceae | 1 | 0 |
| Dothioraceae | 0 | 1 |
| Hyaloscyphaceae | 0 | 1 |
| Meruliaceae | 0 | 1 |
| Pleosporales *incertae*<br>*sedis* | 0 | 1 |
| Sacrosomataceae | 0 | 2 |
| unknown Helotiales | 0 | 1 |
| Xylariaceae | 0 | 1 |

**Table S3.** Order counts for nonserpentine and serpentine fungal communities isolated from *Stipa pulchra* leaves. Numbers in each column represent the number of times an order was isolated from *S. pulchra* leaves in each habitat type.

| **Order** | **Nonserpentine** | **Serpentine** |
| --- | --- | --- |
| Pleosporales | 124 | 65 |
| Capnodiales | 17 | 40 |
| Helotiales | 1 | 3 |
| Sordariales | 1 | 0 |
| Dothideales | 0 | 1 |
| Pezizales | 0 | 2 |
| Polyporales | 0 | 1 |
| Xylariales | 0 | 1 |

**Table S4.** Class counts for nonserpentine and serpentine fungal communities isolated from *Stipa pulchra* leaves. Numbers in each column represent the number of times a class was isolated from *S. pulchra* leaves in each habitat type.

| **Class** | **Nonserpentine** | **Serpentine** |
| --- | --- | --- |
| Dothideomycetes | 141 | 106 |
| Leotiomycetes | 1 | 3 |
| Sordariomycetes | 1 | 1 |
| Agaricomycetes | 0 | 1 |
| Pezizomycetes | 0 | 2 |

**Table S5.** Presence (P, shaded) or absence (A, unshaded) of genera in nonserpentine and serpentine fungal communities isolated from *Stipa pulchra* leaves, by year.

| Genus | Nonserpentine 2015 | Nonserpentine 2016 | Nonserpentine 2017 | Serpentine 2015 | Serpentine 2016 | Serpentine 2017 |
| --- | --- | --- | --- | --- | --- | --- |
| *Alternaria* | P | P | P | P | P | P |
| *Chalastospora* | A | A | A | P | A | A |
| *Cladosporium* | P | A | A | P | A | P |
| *Drechslera* | P | P | P | P | P | P |
| *Epicoccum* | A | A | P | A | A | A |
| *Heyderia* | A | A | P | A | A | P |
| *Hyphodiscus* | A | A | A | A | P | A |
| *Keissleriella* | P | P | P | P | P | P |
| *Nemania* | A | A | A | A | A | P |
| *Neosascochyta* | A | P | P | A | P | P |
| *Parastagonospora* | P | A | P | A | A | A |
| *Phaeophlebiopsis* | A | A | A | A | P | A |
| *Phaeosphaeria* | A | P | A | P | P | A |
| *Plectania* | A | P | A | A | A | P |
| *Podospora* | A | P | A | A | A | A |
| *Pseudoseptoria* | A | A | A | A | P | A |
| *Pyrenophora* | P | P | P | P | P | P |
| *Ramularia* | P | P | A | P | P | P |
| *Stemphylium* | A | P | P | P | A | P |
| unknown<br>Phaeosphaeriaceae | P | P | A | A | P | A |
| unknown<br>Pleosporaceae | A | P | P | A | P | P |
| unknown Helotiales | A | A | A | A | P | A |
| *Wojnowiciella* | A | A | A | P | A | A |

**Table S6.**
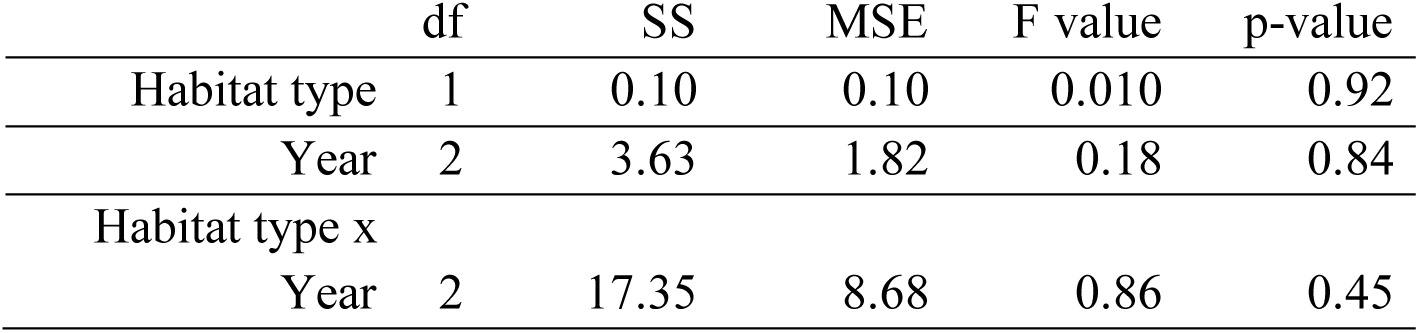
ANOVA table for effects of habitat type, year, and habitat type x year interactions on site-level fungal community diversity in terms of Fisher’s alpha. Columns, from left to right, list the variable tested, the degrees of freedom, the sum of squares, the mean squared error, the F value, and the p-value.

**Table S7.** ANOVA table for effects of habitat type, year, and habitat type x year interactions on site fungal species richness. Columns, from left to right, list the variable tested, the degrees of freedom, the sum of squares, the mean squared error, the F value, and the p-value.

|  | df | SS | MS | F value | p-value |
| --- | --- | --- | --- | --- | --- |
| Habitat type | 1 | 6.72 | 6.72 | 1.07 | 0.32 |
| Year | 2 | 3.11 | 1.56 | 0.25 | 0.78 |
| Habitat type x<br>year | 2 | 8.44 | 4.22 | 0.67 | 0.53 |

**Table S8.**
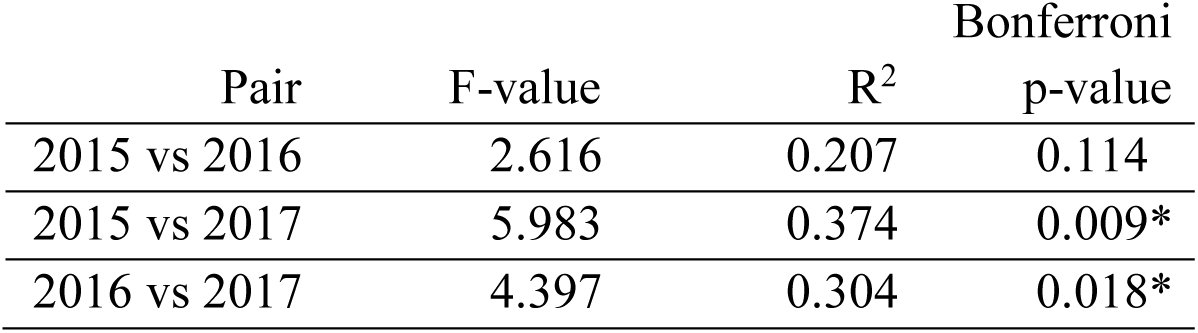
Pairwise PERMANOVA table for effects of year on fungal community composition. From left to right, columns list the pair of years being compared, the F-value, R^2^, and the Bonferroni-adjusted p-value. P-values < 0.05 are marked with an asterisk.

**Table S9.** Results of chemical analysis of serpentine and nonserpentine *S. pulchra* foliar tissue. The Serpentine and Nonserpentine columns report elemental content in percentage of leaf tissue by dried weight. The standard error of each mean is listed in parentheses. W, in the third column, is the Wilcoxon-rank sum test statistic comparing serpentine and nonserpentine mean values. Asterisks denote statistically significant p-values (p < 0.05).

| Element | Serpentine |  | Nonserpentine |  | W | p-value |  |
| --- | --- | --- | --- | --- | --- | --- | --- |
| C | 38.97 | $(3.44 \times 10^{-1})$ | 41.48 | $(2.35 \times 10^{-1})$ | 2713 | $1.65 \times 10^{-7}$ | * |
| N | 1.44 | $(4.96 \times 10^{-2})$ | 1.61 | $(3.84 \times 10^{-2})$ | 2346 | $1.10 \times 10^{-3}$ | * |
| C:N | 28.72 | $(8.74 \times 10^{-1})$ | 26.54 | $(6.18 \times 10^{-1})$ | 1398 | $6.62 \times 10^{-2}$ | |

**Table S10.** Analysis of deviance table for multivariate test of fungal community responses, based on fungal species presence/absence data, to foliar percent C, foliar percent N, foliar C:N ratio, and habitat type. Only fungal species represented by five or more isolates in 2016 were considered.

|  | df | deviance | p-value |
| --- | --- | --- | --- |
| Habitat type | 96 | 20.93 | 0.01* |
| Foliar %C | 95 | 7.98 | 0.35 |
| Foliar %N | 94 | 9.02 | 0.33 |
| Foliar C:N | 93 | 6.896 | 0.63 |

